# Epigenomic Landscape of *Arabidopsis thaliana* Metabolism Reveals Bivalent Chromatin on Specialized Metabolic Genes

**DOI:** 10.1101/589036

**Authors:** Kangmei Zhao, Seung Y. Rhee

## Abstract

Metabolism underpins development and physiology, but little is known about how metabolic genes and pathways are regulated, especially in multicellular organisms. Here, we identified regulatory patterns of 16 epigenetic modifications across metabolism in *Arabidopsis thaliana*. Surprisingly, specialized metabolic genes, often involved in defense, were predominantly regulated by two modifications that have opposite effects on gene expression, H3K27me3 (repression) and H3K18ac (activation). Using camalexin biosynthesis genes as an example, we confirmed that these two modifications were co-localized to form bivalent chromatin. Mutants defective in H3K27m3 and H3K18ac modifications showed that both modifications are required to determine the normal transcriptional kinetics of these genes upon stress stimuli. Our study suggests that this type of bivalent chromatin, which we name a kairostat, controls the precise timing of gene expression upon stimuli.

**One Sentence Summary:** This study identified a novel regulatory mechanism controlling specialized metabolism in *Arabidopsis thaliana*.

---

Plants synthesize a plethora of compounds, many of which are used as food, feed, fuel, and pharmaceuticals (*1, 2*). Genes responsible for the biosynthesis of these compounds are wired into a complex network. Various levels of regulation, including epigenetic modifications and transcription factors, play important roles in coordinating the expression of genes. However, compared to developmental processes, knowledge about the epigenetic regulation of metabolism is limited in multicellular organisms. In Arabidopsis, trimethylation of lysine 36 in histone 3 (H3K36me3) activates energy related pathways (*3*) and trimethylation of lysine 27 in histone 3 (H3K27me3) represses lipid biosynthesis pathways (*4*). However, the genome-wide epigenetic regulatory landscape of metabolism in plants is not known.

To determine how metabolism is regulated epigenetically, we first identified regulatory patterns of epigenetic modification across all metabolic domains. Specifically, we examined 16 high-resolution epigenomic profiles, including histone variants, DNA methylation, and histone modifications (*5*). To map the epigenetic marks on metabolic enzymes, pathways, and domains, we used the genome-wide functional annotations of metabolism we generated previously (*6*). To see if there are any interactions between epigenetic modifications across metabolic genes, we computed pairwise Pearson’s correlation coefficients between the 16 epigenetic modifications based on their relative abundance at each metabolic gene region. Four groups of epigenetic regulons were identified (Fig. 1A). Groups I and II included mainly the epigenetic modifications that activate gene expression. Group III included only the modifications that repress expression, such as H3K9me2 and H3K27me1. Interestingly, group IV included two modifications that have opposite effects on expression: H3K18ac, an activation mark, and H3K27me3, a repression mark. We further asked how different domains of metabolism are regulated by epigenetic modifications. Enrichment of modifications in each metabolic domain, relative to all the genes in the genome, revealed both shared and diverse patterns. For example, energy, amino acid, and nucleotide metabolic domains, which are essential for survival, growth, and development, were significantly enriched with marks that activate gene expression, such as H3K36me2, H3K36me3 H2Bub, H3K2me2, H3K4me3, and H3K9ac (Fisher’s exact test, p value < 0.05, fold change ≥ 1.5) (Fig. 1B). In contrast, metabolic genes involved in specialized and hormone metabolic domains, which are generally involved in interactions with biotic and abiotic environmental stimuli, showed distinct patterns with the enrichment of a repression mark H3K27me3 and an activation mark H3K18ac (Fig. 1B). The enrichment patterns of epigenetic modifications were consistent with expression levels of genes involved in each domain under healthy conditions (Fig. S1A). For example, genes associated with energy metabolism showed higher expression levels compared to those involved in specialized metabolism (Fig. S1B).

**Fig. 1.**
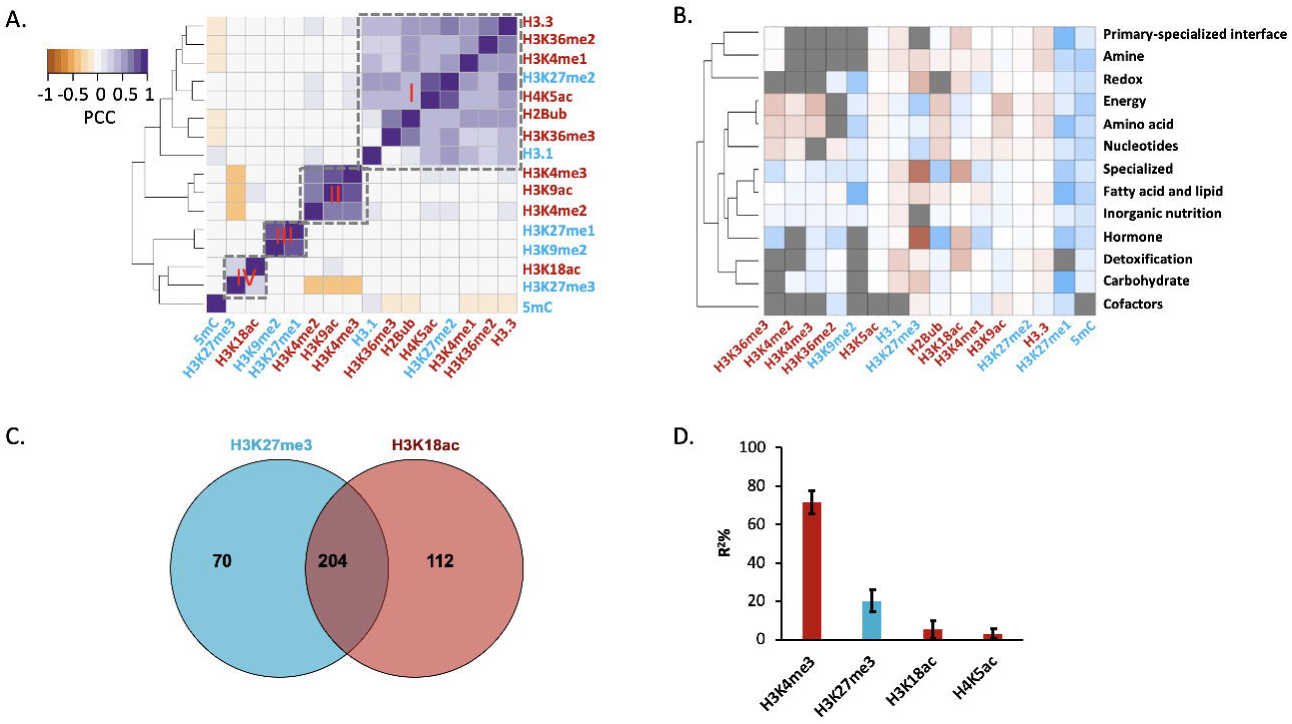
Epigenetic modifications may functionally cooperate to regulate metabolism. A. Correlation analysis reveals four groups of epigenetic marks, activating (red) and repressing (blue), based on their relative abundance on metabolic genes. B. Enrichment analysis shows differential epigenetic modification patterns across metabolic domains. The heatmap represents log2 fold change of enrichment or depletion of a mark relative to all genes in the genome based on Fisher’s exact test. Gray cells represent non-significant comparison. C. The association between H3K27me3 and H3K18ac on specialized metabolic genes is higher than expected (hypergeometric test, p value = 0.03). D. Multiple linear regression and bootstrap-based variable selection show that the absence of H3K4me3 and presence of H3K27me3 are the most predominant factors that determined the expression of specialized metabolic genes.

To understand how prevalent these observed epigenetic modification patterns were for each metabolic pathway, we performed pathway-level enrichment analysis. Pathways with fewer than 10 genes were excluded from the analysis to avoid the small sample size problem. For energy metabolism, more than 50% of the pathways were significantly enriched with activation marks, including H3K4me2, H3K4me3, and H3K36me3 (Fig. S2). Moreover, 52% (168 out of 321) of energy-related metabolic genes were targets of all these three modifications (Fig. S2B). On the other hand, repression marks such as DNA 5’ cytosine methylation and H3K27me3 were significantly depleted across energy-metabolism related pathways (Fig. S2A). These patterns suggest that multiple epigenetic modifications that activate expression maintain the high expression level of energy-related metabolism genes, though we cannot distinguish the marks that work together on the same genes from those that harbor different modifications on the same genes in different cell types. For specialized metabolism, 74% (39 out of 53) of the pathways were enriched with both H3K18ac and H3K27me3 (Fig. S3). Genes involved in specialized metabolism were more likely to be regulated by both modifications than expected by chance in the genome (hypergeometric test, p value = 0.03, fold change = 1.3) (Fig. 1C). This suggests that H3K18ac and H3K27me3 may work together somehow to regulate the expression of specialized metabolic genes. Multiple linear regression models further confirmed that the significantly enriched and depleted epigenetic modifications predominantly determine the expression patterns of energy and specialized metabolic genes (Figs. 1D, S4, S5, and S6). Interestingly, the absence of H3K4me3 explained about 70% of the expression level of specialized metabolic genes and the presence of H3K27me3 and H3K18ac explained 27% of the expression (Fig. 1D). This suggests that a few epigenetic marks are sufficient to predict gene expression levels.

Epigenomic patterns suggested that H3K18ac and H3K27me3 could be important for regulating the expression of genes involved in specialized metabolism. Moreover, gene targets of these two modifications overlapped more than expected by chance. This may be explained by two mutually non-exclusive hypotheses: 1) H3K18ac and H3K27me3 are co-localized on the same genes involved in specialized metabolism in the same cell; or 2) H3K18ac and H3K27me3 regulate the expression of the same genes but in different cells, which may contribute to determining the cell-type specificity of specialized metabolism. To distinguish between the two possibilities and understand the role of H3K27me3 and H3K18ac on regulating gene expression, we examined *in vivo* co-localization of the two modifications using sequential chromatin immunoprecipitation (ChIP)-qPCR, which requires a two-step, serial chromatin pull-down with antibodies against these two modifications. We selected the camalexin biosynthesis pathway as an example representing specialized metabolism. Camalexin is a simple indole alkaloid derived from tryptophan and is one of the most potent defense compounds against bacterial and fungal infections in Arabidopsis (*7*). Major genes encoding enzymes that catalyze each reaction in this pathway have been functionally characterized, including *CYP79B2, CYP71A12*, and *PAD3* (*8, 9*) (Fig. 2A). To assess the antibodies’ specificity and efficiency, we included various positive and negative controls in the ChIP assay and tested a transcription factor encoding gene called *Golden-2-Like 1* (*GLK*) that was previously observed to be modified by both H3K27me3 and H3K18ac (*10*). All three camalexin biosynthesis genes showed significantly higher signals when pulled down with both antibodies against H3K27me3 and H3K18ac than without any antibody or with the same antibody in the second pull-down (Fig. 2B). We observed comparable pull-down efficiency for GLK. Altering the order of the two antibodies for H3K27me3 and H3K18ac in the pull-down showed similar results (Fig. 2C). These results indicated that H3K18ac and H3K27me3 are co-localized at the camalexin biosynthesis genes *in planta* to form bivalent chromatin.

**Fig. 2.**
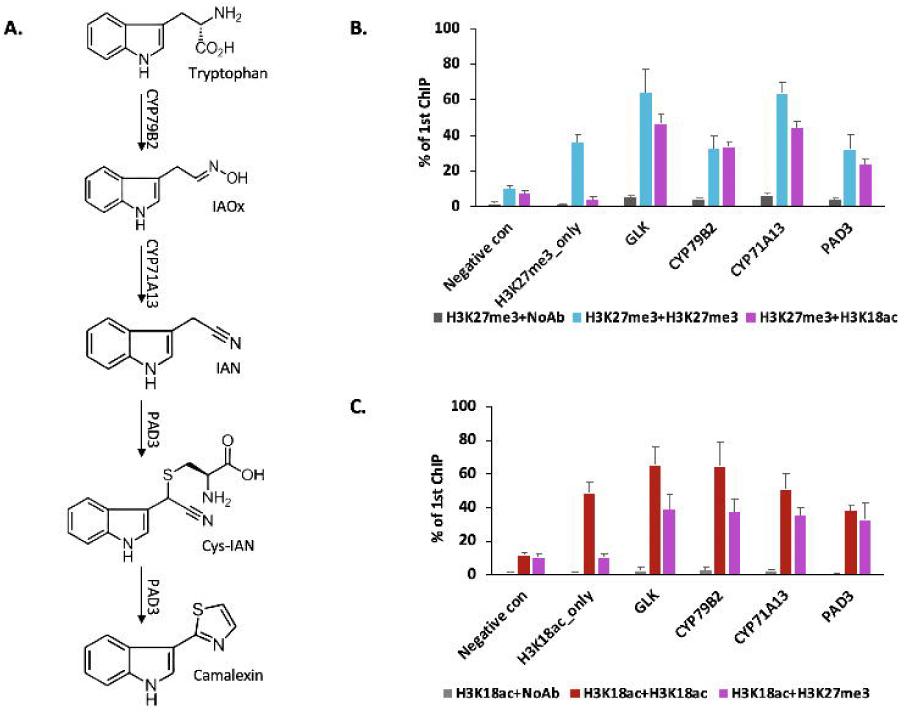
H3K27me3 and H3K18ac are co-localized on camalexin biosynthesis genes to form bivalent chromatin. A. Camalexin biosynthesis pathway and the minimum set of genes that can produce the final compound. B Sequential ChIP-qPCR confirms the co-localization of H3K27me3 and H3K18ac. C. Altering the order of antibodies shows similar results in sequential ChIP-qPCR. Negative control represents a genomic region that has very low abundance for H3K27me3 and H3K18ac based on the publicly available ChIP-seq data (5). H3K27me3_only represents a genomic region that has very low abundance of H3K27me3 (5). H3K18ac_only represents a genomic region that has very low H3K18ac. Error bars represent standard deviation.

Bivalent chromatin has been observed in gene regions associated with human stem cell and cancer cell development, as well as a few other loci in animal and Arabidopsis genomes, including the *FLOWERING LOCUS C* (*10-14*). The biological function of bivalent chromatin has been hypothesized to poise gene expression for rapid activation upon signaling. To date, no direct evidence is available to test this hypothesis in a whole organism context. To understand the biological function of bivalent chromatin on the regulation of specialized metabolism, we examined the transcriptional kinetics of camalexin biosynthesis genes under Flagellin 22 (FLG22) induction in wild type as well as mutant lines that are significantly reduced in the deposition of H3K27me3, *pkl-1(15)*, and H3K18ac, *hda18-1* (*16*). First, we confirmed that the abundance of H3K27me3 and H3K18ac at the loci of camalexin biosynthesis genes was significantly reduced in *pkl-1* and *hda18-1* mutant plants compared to the wild type using ChIP-qPCR (Figs. S7, S8). For each gene tested in the experiment, primers were designed to cover the region containing 1kb upstream and the entire transcribed region (Table S3). We then examined the transcriptional kinetics of genes in the camalexin biosynthesis pathway in response to FLG22 induction. Under control conditions, the expression levels of all three genes were similar in *pkl-1* or *hda18-1* lines compared to those in wild type plants (Fig. 3). This indicates that bivalent chromatin may not be needed to maintain the low expression level of specialized metabolic genes under healthy conditions. However, with FLG22 treatment, *CYP79B2* was induced within 5 minutes after the treatment. The other two genes within this pathway, *CYP71A13* and *PAD3*, showed a dramatic expression change 45 minutes after the treatment in wild type plants (Fig. 3). For *pkl-1* lines (reduced H3K27me3 marks), *CYP71A13* and *PAD3* showed a significant induction of gene expression within 5 minutes of the treatment and the degree of induction was much higher than that in wild type plants (Fig. 3). In *hda18-1* mutants (reduced H3K18ac marks), for all three genes, a significant induction of expression occurred around 60 minutes after FLG22 treatment (Fig. 3). These results demonstrate that H3K18ac-H3K27me3 bivalent chromatin controls the timing of expression of camalexin biosynthesis genes upon stress signal, with H3K18ac expediting and H3K27me3 attenuating expression to hit the presumed “sweet spot” (Fig. 4). We have named this type of bivalent chromatin regulator a kairostat, inspired by the ancient Greek word ‘kairos’ meaning the right moment and ‘stat’ meaning regulating device.

**Fig. 3.**
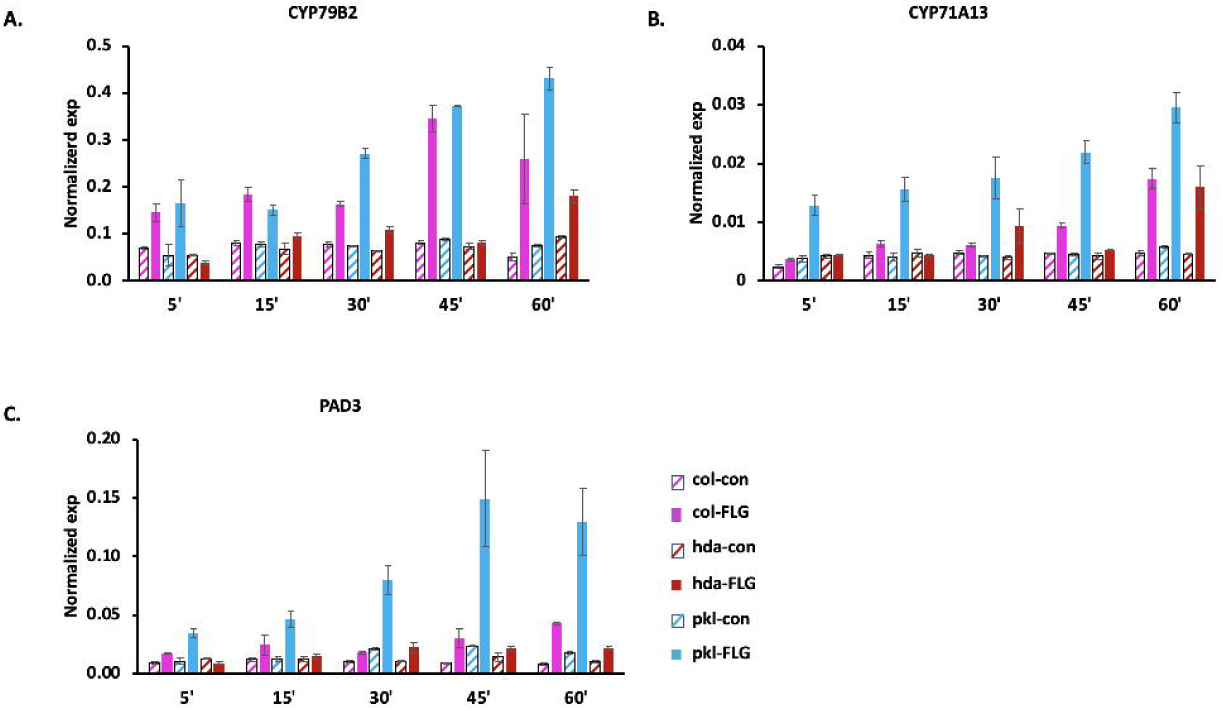
Transcriptional kinetics analysis shows that the H3K27me3-H3K18ac bivalent chromatin does not affect the expression of camalexin biosynthesis genes under healthy conditions, but controls the timing of gene expression induction upon stress signals. A, B, and C represent the expression of the three major genes in camalexin biosynthesis pathway. ACT2 was used as the reference gene in this experiment. Two-week old Arabidopsis seedlings were treated with 1µm FLG22 in this experiment. Error bars represent standard deviation.

**Fig. 4.**
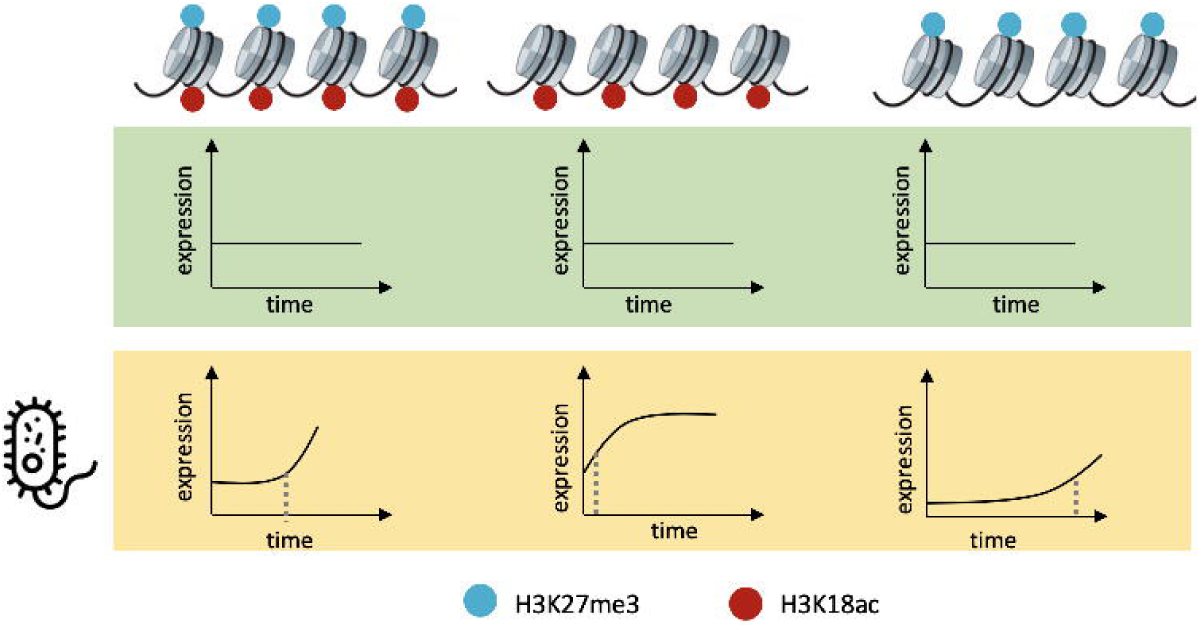
A model of a bivalent chromatin that acts as a kairostat on regulating the expression of some specialized metabolic genes. Under healthy conditions, bivalent chromatin does not seem to control gene expression. Under FLG22 treatment, bivalent chromatin controls the timing of camalexin biosynthesis genes, with H3K18ac expediting and H3K27me3 attenuating expression to hit the presumed ‘sweet spot’. Dashed lines represent the time of gene induction

In this study, we systematically identified novel regulatory patterns of metabolism at the epigenetic level in a whole organism by integrating omics data with genetics. Metabolic genes that are essential for survival, such as those involved in energy metabolism, show active expression, which is maintained by the enrichment of activation marks and depletion of repression marks. In contrast, our data suggests that specialized metabolism is regulated partly by bivalent chromatin, formed by the colocalization of H3K18ac and H3K27me3. Bivalent chromatin has so far only been characterized in human stem and cancer cell models, and observed in a few loci of animal and plant models *(10-14, 17)*. In this study, we functionally examined the role of bivalent chromatin on gene expression regulation upon stress induction of Arabidopsis plants. Interestingly, the transcriptional kinetics of camalexin biosynthesis genes suggests that the bivalent chromatin does not affect expression under healthy conditions, but rather controls the timing of gene expression upon a stimulus. We have named this type of regulator a kairostat. Elucidating how kairostats function could have far-reaching implications not only in basic and synthetic biology, but also in other fields such as agriculture, engineering, and medicine.

## Acknowledgments

We thank Jennifer Brophy and Flavia Bossi for their critical feedback on this work. We acknowledge Benjamin Jin for help with identifying homozygous mutant lines for *hda18-1*. We also acknowledge Jia-Ying Zhu and Yuchun Hsiao for help with ChIP-qPCR experiments and Hye-In Nam for help with growing plants. **Funding**: This work was supported in part by Carnegie Institution for Science Endowment and grants from the National Science Foundation (IOS-1546838, IOS-1026003), the Department of Energy (DE-SC0008769, DE-SC0018277), and the National Institutes of Health (1U01GM110699-01A1). **Author contributions**: K.Z. conducted computational analysis, ChIP, and transcriptional kinetics experiments. S.R. supervised this study. K.Z. and S.R. wrote the manuscript. **Competing interests:** The authors claim no competing interests. **Data and materials availability**: All data are available in the manuscript or in the supplementary materials.

## Supplementary Materials

Materials and Methods Figures S1-S7

Tables S1-S5

---

**Data S1.** Log2(fold change) in the enrichment analysis to identify predominant epigenetic modifications in energy metabolism-related pathways. Only the pathways that contain at least ten genes were included in this analysis.

**Data S2.** Log2(fold change) in the enrichment analysis to identify predominant epigenetic modifications in specialized metabolism-related pathways. Only the pathways that contain at least ten genes were included in this analysis.

