## Supplementary Material for "Epigenomic Landscape of *Arabidopsis thaliana* Metabolism Reveals Bivalent Chromatin on Specialized Metabolic Genes"

**This PDF file includes:**

Materials and Methods
Figs. S1 to S8
Tables S1 to S3

**Other Supplementary Materials for this manuscript include the following:**

Data S1 to S2

### Materials and Methods

#### Annotation of Metabolic Genes, Pathways, and Domains

Arabidopsis metabolic enzymes and pathways were extracted from Plant Metabolic Network (PMN) (http://www.plantcyc.org/). Enzymes were annotated based on the Ensemble Enzyme Prediction Pipeline (E2P2) (*6, 18*), metabolic pathways were predicted by Pathway Tools software (*19*), which were further validated by a process called Semi-Automated Validation and Integration (SAVI) (*6, 20*). In *A. thaliana*, 5451 genes were predicted to encode enzymes that catalyze 3585 reactions in small molecule metabolism, which were assigned to 627 pathways based on prediction and manual curation based on PlantCyc version 12 (*6*). These pathways were further grouped into 13 metabolic domains based on the types of metabolites that they potentially synthesize. The metabolic domain(s) assigned to a given pathway were transferred to the reactions, EC numbers, and proteins/genes associated with that pathway. To improve the power of statistics, we included only those pathways that contain more than ten genes when we performed epigenetic modification enrichment analysis on individual pathways in energy or specialized metabolism.

#### Expression Patterns of Metabolic Domains

To analyze the expression level of genes associated with various metabolic domains under healthy conditions, we used the transcriptomic dataset generated using 10-day old *Arabidopsis thaliana* seedlings grown on 0.5X MS medium containing 1% sucrose at 23°C under long days (16 h light/8 h dark) (*5*). This is a comparable condition as the experiments conducted in our study. The RNA-seq reads were mapped to *Arabidopsis thaliana* genome v.10 using the STAR package with default settings (*21*). After the reads were mapped, gene expression level represented by Reads Per Kilobase of transcript per Million (RPKM) was calculated with the GenomicRange R package v3.5 (*22*). To determine the level of expression for each gene, the RPKM values were further ranked and assigned an integer score (1,9). These gene expression levels were used in the expression pattern analysis and multiple linear regression models.

#### Epigenomic Enrichment Analysis

The 16 epigenomic profiles were downloaded from Wang et al 2015, where previously published ChIP-seq and ChIP-chip data for all epigenetic marks were re-mapped to *Arabidopsis thaliana* genome v.10 (*5*). Briefly, for each ChIP-seq profile, the signals were binned and ranked in the genome (*5*). We used the ranked epigenetic modification signals in the following analyses.

To identify the predominant epigenetic modifications associated with metabolic domains and pathways, we first mapped the ranked epigenetic modification signals to each gene, including the transcribed region, 1kb upstream, and 500bp downstream regions. Then, we calculated the average epigenetic modification density for each domain or pathway by taking the sum of the ranked modification signals observed within all the genes in that domain or pathway and normalizing it to the total length of all the genes. To identify enriched marks within metabolic domains or pathways, we compared the density of each epigenetic modification within that domain or pathway to the background signal. We defined the background signal as the average density of each epigenetic modification by taking the sum of the epigenetic modification signals observed within total gene regions in the genome and normalizing it to the length of total genes. Fold change of enriched or depleted epigenetic modifications for energy and specialized metabolic pathways were summarized in Data S1 and S2. The statistical significance of enrichment or depletion was determined by a G test and followed by a post-hoc adjustment using FDR with the threshold of 0.01. The heat maps were generated with gplots v.3 (*22*) and ggplot2 version 3.1 (*23*) packages in R.

**Multiple Linear Regression to Predict Expression Based on Epigenomic Datasets**

Multiple linear regression was applied to model gene expression levels based on 16 epigenomic profiles of all metabolic genes. We randomly split total metabolic genes into training and validation sets with the same number of variables and applied best model selection with various number of epigenetic marks using an R package called ISLR version1.2 (*24*). Cross validation returned an eight-variable model as the best model based on adjusted $R^2$ and Bayesian Information Criterion (BIC). BIC represents model complexity and, as long as the value of the BIC decreases, increasing the model complexity can improve prediction power. We assessed the prediction power of the best model by determining the Pearson's correlation coefficient between predicted expression using the best full model and measured expression using publicly available RNA-seq data described above. The result showed that the model computed with total metabolic genes explained 61% of expression levels ($r^2 = 0.61$; Fig S2C), which showed a similar prediction power compared to the models built using epigenomic datasets in human cells and *Caenorhabditis elegans* (*25*). We then asked whether we could identify the most informative epigenetic marks that were able to explain the expression of metabolic genes involved in energy or specialized metabolism. Based on model fitness measured by adjusted $R^2$ and model complexity reflected by BIC, we identified the best models containing five variables to infer the expression of energy metabolism and four variables for specialized metabolism. To further identify the relative importance of individual modification on the prediction of expression within these two best models, we applied a bootstrap-based algorithm using the R package RELAIMPO (*26*). The bootstrapping was done by randomly sampling the residuals of the regression model. The energy metabolism model revealed five informative variables for the prediction of expression (model fitness: $r^2 = 0.45$, F-test p value < 2E-16): H3K4me3, H3K27me3, H3K9me2, H4K5ac, and H3K36me3. The specialized metabolism model revealed four informative variables for the prediction of expression (model fitness $r^2 = 0.51$, F-test, p value < 2E-16): H3K4me3, H3K18ac, H3K27me3, and H4K5ac.

**Plant Growth Conditions**

Seeds of *Arabidopsis thaliana* accession Columbia (Col-0) were stratified at 4°C for three days before germination. Then plants were grown in 0.5X MS medium containing 1% sucrose and 0.8% agar at 23°C under long days (16 h light/8 h dark) with controlled light at 100 µmol photosynthetic photons/$m^2$ s (PPFD). Two-week old seedlings were harvested for the ChIP-qPCR, sequential ChIP-PCR, and transcriptional kinetics experiments.

**ChIP-qPCR and Sequential ChIP-PCR**

Seedlings were cross-linked with 1% formaldehyde by vacuum infiltration and quenched with 0.125 M glycine. After grinding, nuclei were isolated in an extraction buffer containing 10 mM Tris-HCl pH 8.0, 0.25 M sucrose, 10 mM $MgCl_2$, 1% Triton X-100, and protease inhibitors. Nuclei were pelleted by centrifugation, resuspended in lysis buffer (50 mM Tris-HCl, pH 8.0, 10 mM EDTA, and 1% SDS) and disrupted by sonication in a Bioruptor (Diagenode), yielding genomic fragments of 200 to 600bp.

To confirm that the mutant lines, *hda18-1* and *pkl-1*, have reduced histone modifications, ChIP-qPCR was used to quantify the abundance of H3K27me3 and H3K18ac through the locus of all three camalexin biosynthesis genes. After chromatin crosslinking and shearing, DNA/protein complexes were immunoprecipitated with antibodies against H3K27me3 (Milipore, Cat No 07-449) and H3K18ac (Abcam ab8580) by referring the ChIP protocol established based on Arabidopsis tissues (*27*). DNA was purified using QIAquick PCR Purification Kit (Qiagen, Cat No 28104). Primers were designed to cover the 1kb upstream of the transcription start site and the entire transcribed region (Table S1).

Sequential ChIP experiments were performed to examine the co-localization of H3K27me3 and H3K18ac on camalexin biosynthesis genes. Chromatin crosslinking and shearing followed the protocol described above for ChIP-qPCR. For each ChIP/re-ChIP assay, 8 to 15 µg of DNA/protein complexes were immunoprecipitated with anti-H3K27me3 (Milipore, Cat No 07-449) and anti-H3K18ac (Abcam ab8580) antibodies using the Re-ChIP-IT kit (Active Motif, Cat No 53016). Quantitative real-time PCR was performed using Roche 480 detecting system with the SensiFAST Sybr No-Rox mix (Bioline, Cat No BIO98020) from Bioline. Primer sequences used in this study are provided in Table S2. To represent final results after two-step chromatin immunoprecipitation, the pull-down efficiency was calculated as the percent of first ChIP using the equation: $1^{st}$ ChIP = $2^{\wedge(Cq\_2nd\ ChIP\ -\ Cq\_1st\ ChIP)}$.

**Transcriptional Kinetics Analysis**

Two-week old seedlings grown in the condition as described above were used to study the transcriptional kinetics. To induce the expression of camalexin biosynthesis genes, seedlings were treated with Flagelin22, a short peptide synthesized at Stanford PAN Center, with 1µM final concentration. Then tissues were sampled at 5, 15, 30, 45, and 60 minutes after FLG22 treatment and frozen immediately in liquid nitrogen.

After tissue grinding, RNA was extracted using RNeasy Plant mini kit (Qiagen, Cat No 74904) and 1µg RNA was used for cDNA synthesis using SuperScript First-Strand Synthesis System (Thermo Fisher, Cat No 18080051). Quantitative real-time PCR was performed using Roche 480 detecting system with the SensiFAST Sybr No-Rox mix (Bioline, Cat No BIO98020) from Bioline. Primers are summarized in Table S3. One of the housekeeping genes, Actin 2 (AT3G18780), was used as the reference gene in the qPCR experiment.

A.

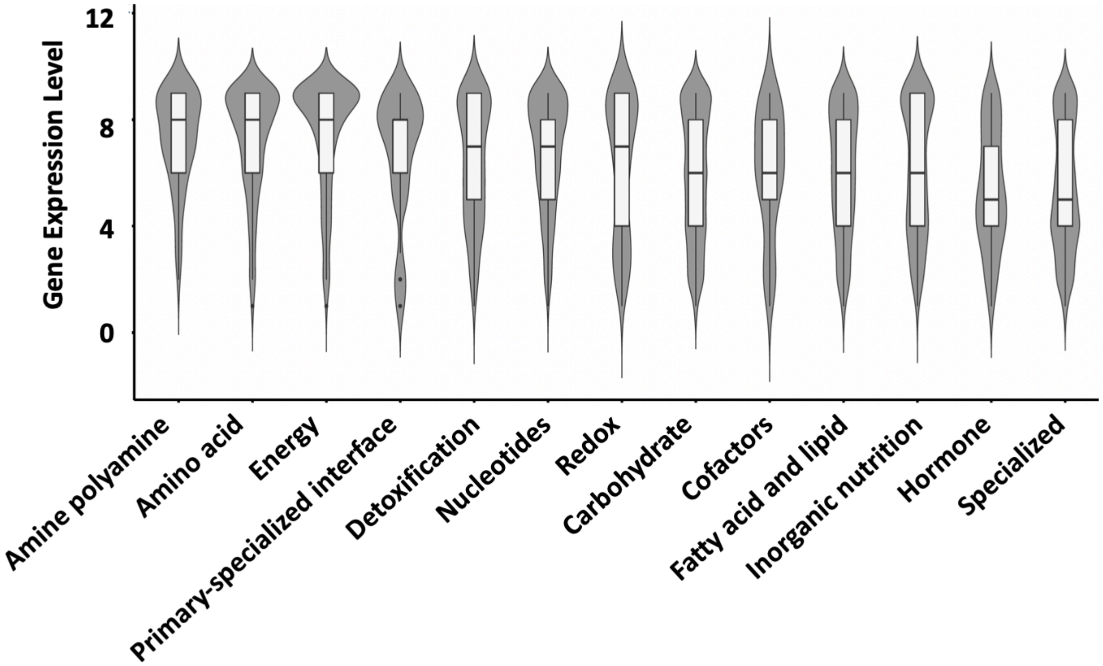


B.

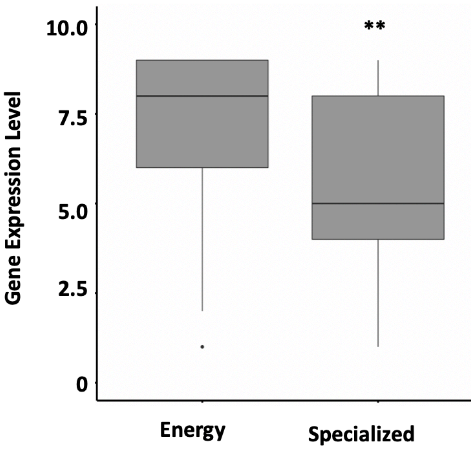


Figure S1. Metabolic domains show various expression levels. A. The expression levels of genes associated with each metabolic domain under healthy conditions. B. Genes involved in energy metabolism show significantly higher expression levels than those associated with specialized metabolism. The RNA-seq data for gene expression level analysis was extracted from NCBI SRA database under the accession number SRP032990, which was generated using ten-day old Arabidopsis seedlings under healthy condition (*5*). Violin and box plots were generated using the package ggplot2 version 3.1 and the domains were sorted based on the median of expression levels in each metabolic domain. ** represents Student's two-tailed T test, p value < 0.01.

A.

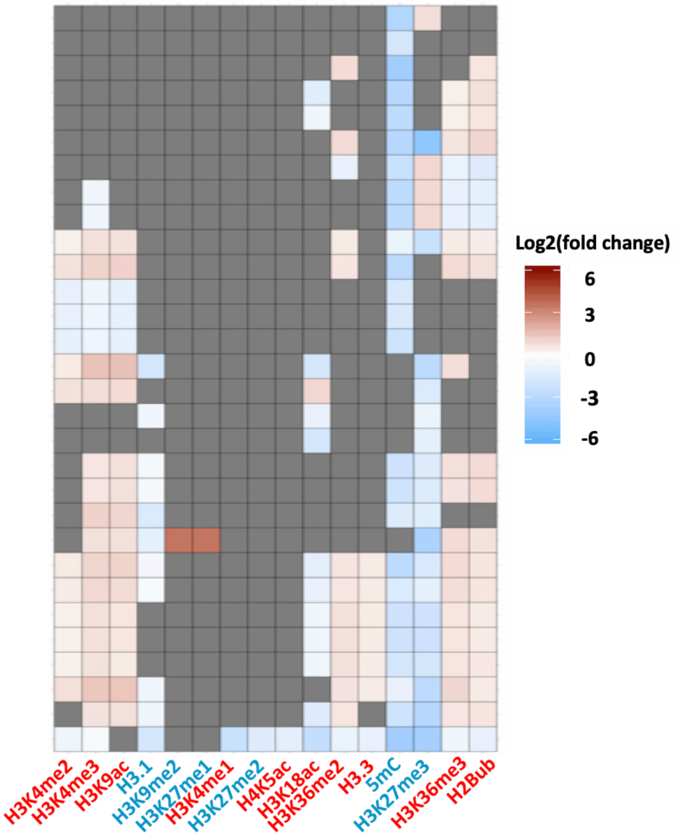


B.

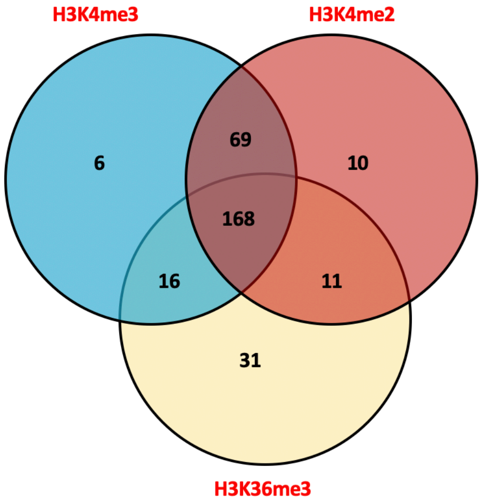


Figure S2. Predominant epigenetic marks associated with energy metabolism-related pathways and genes. A. Enrichment analysis reveals predominant epigenetic modifications in energy metabolism-related pathways. Only the pathways that contain at least ten genes were included in this analysis. The heatmap represents log2 fold change of enrichment or depletion of a mark relative to all genes in the genome based on Fisher's exact test. Gray cells represent non-significant comparison. The heatmap was generated using ggplot2 package version 3.1 in R with hierarchical clustering. Pathway names and fold change for the enrichment analysis are summarized in Data S1. B. Overlapping target genes of H3K4me2, H3K4me3, and H3K36me3 marks suggests putative functional cooperation between these marks on the regulation of energy metabolism-related genes. Epigenetic modifications were color-coded based on their effect on gene expression; red represents activation marks and blue represents repression marks.

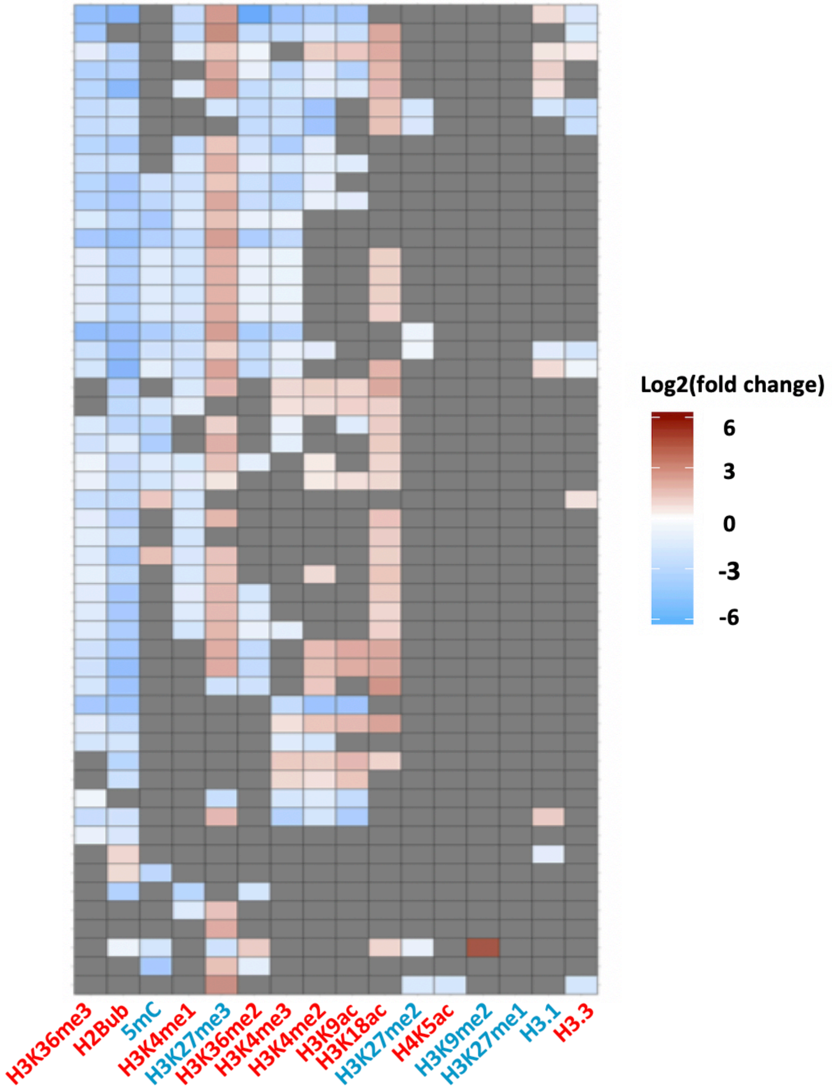


Figure S3. The two prominent epigenetic modifications, H3K27me3 and H3K18ac, are enriched in 74% of specialized metabolism-related pathways. The heatmap represents log2 fold change of enrichment or depletion of a mark relative to all genes in the genome based on Fisher's exact test. Gray cells represent non-significant comparison. The heatmap was generated using hierarchical clustering using the ggplot2 package version 3.1 in R. Gray represents non-significant change based on G test comparing to the background defined by average epigenetic mark density within total gene coding regions. Pathway names and fold change for the enrichment analysis are summarized in Data S2. Epigenetic modifications are color-coded based on their effect on gene expression; red represents activation marks and blue represents repression marks.

A.

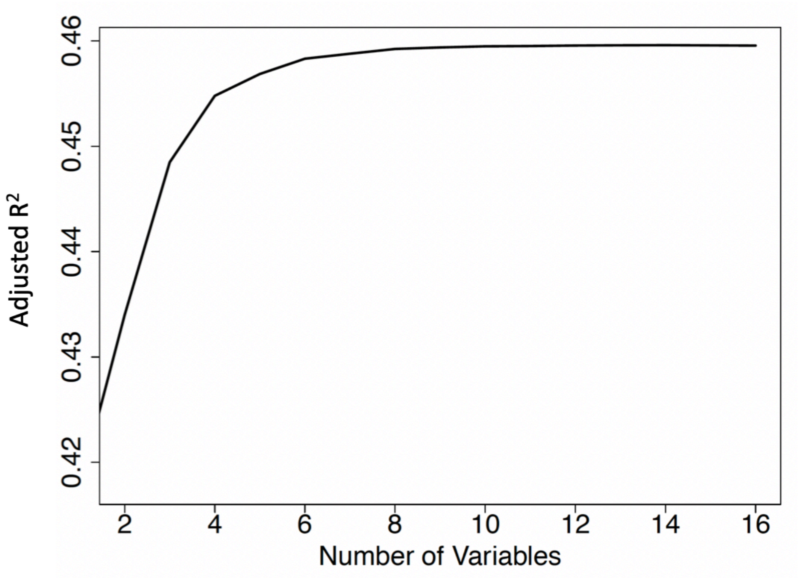
0.46
0.45
0.44
0.43
0.42
Adjusted R2
2
4
6
8
10
12
14
16
Number of Variables

B.

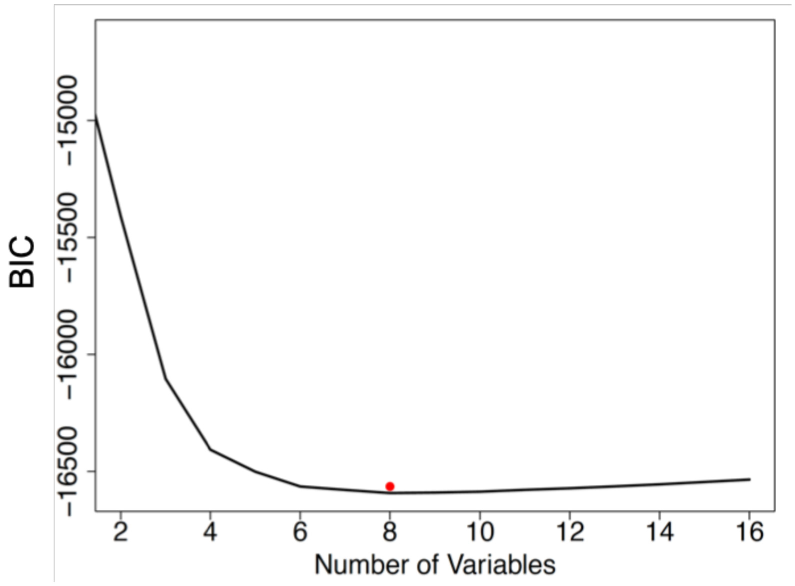
−15000
−15500
−16000
−16500
BIC
2
4
6
8
10
12
14
16
Number of Variables

C.

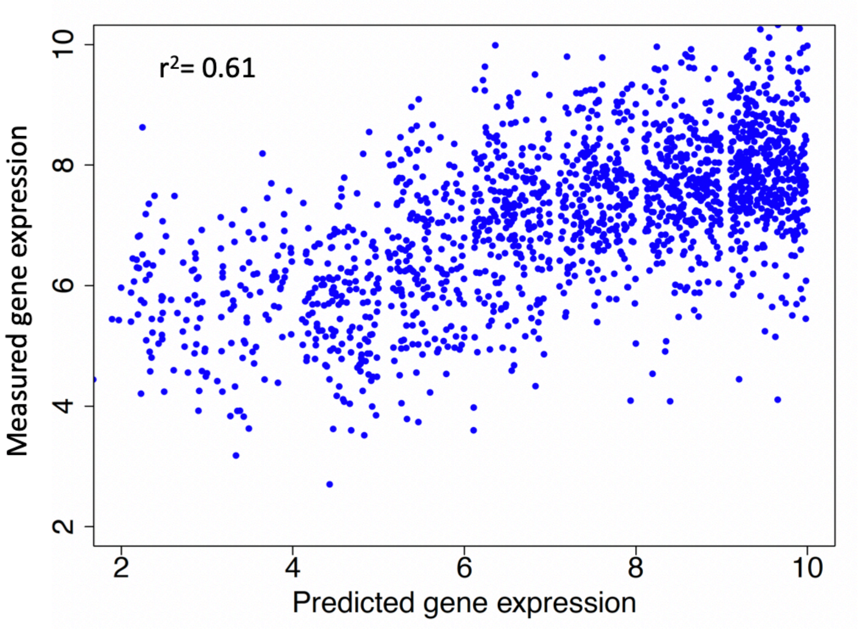


Figure S4. Multiple linear regression shows epigenetic modifications can predict gene expression levels and the best model containing 8 variables explained 61% of expression for total metabolic genes. A. The performance of models generated by different combinations of epigenetic marks. B. Bayesian Information Criterion (BIC) reveals the relationship between model complexity and prediction accuracy and as long as BIC drops, introducing additional epigenetic marks will improve model performance. The red dot represents the number of variables to achieve the least complex and most accurate model. C. The scatterplot shows the expression levels predicted by the best model containing 8 variables on the x-axis and measured values by RNA-seq on the y-axis for total metabolic genes. $r^2$ represents Pearson correlation coefficient between predicted and experimentally measured gene expression levels.

A.

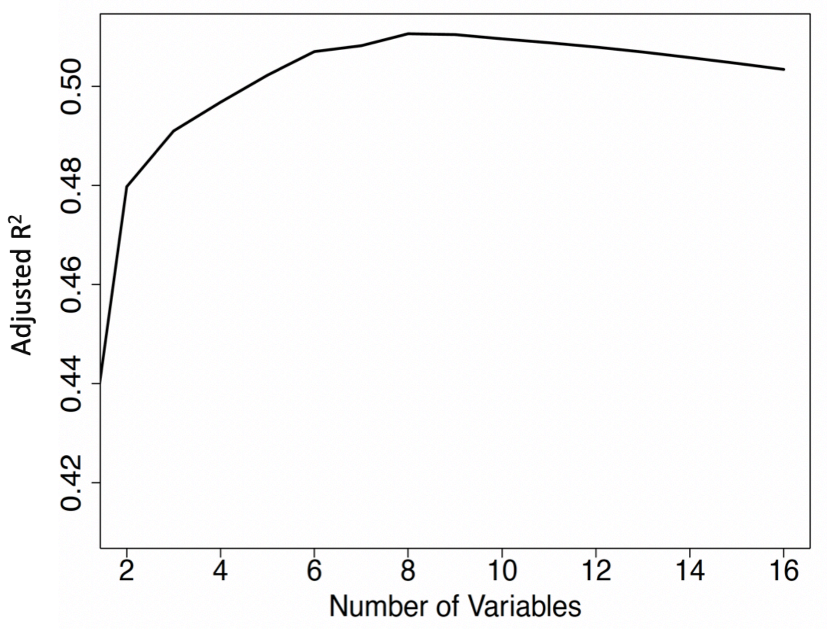
Adjusted R²
0.42
0.44
0.46
0.48
0.50
2
4
6
8
10
12
14
16
Number of Variables

B.

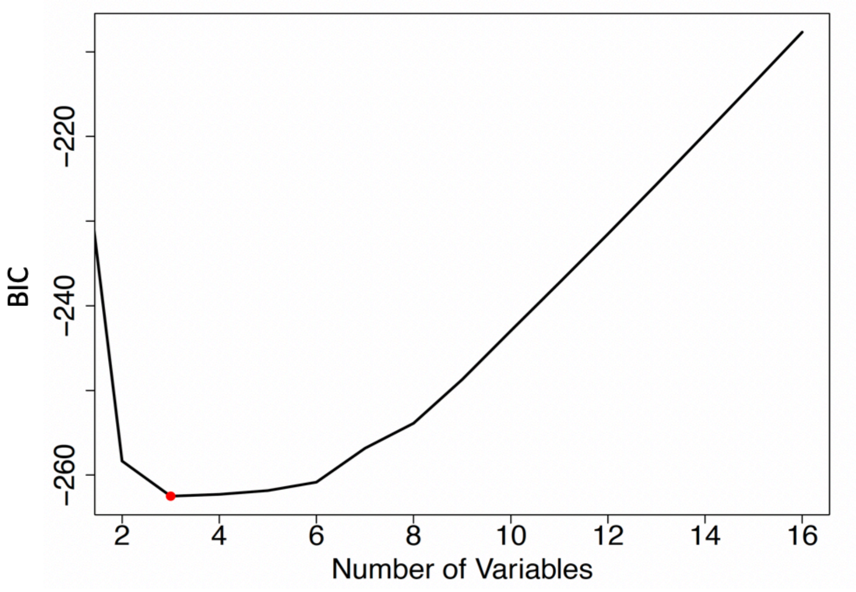
BIC
−260
−240
−220
2
4
6
8
10
12
14
16
Number of Variables

C.

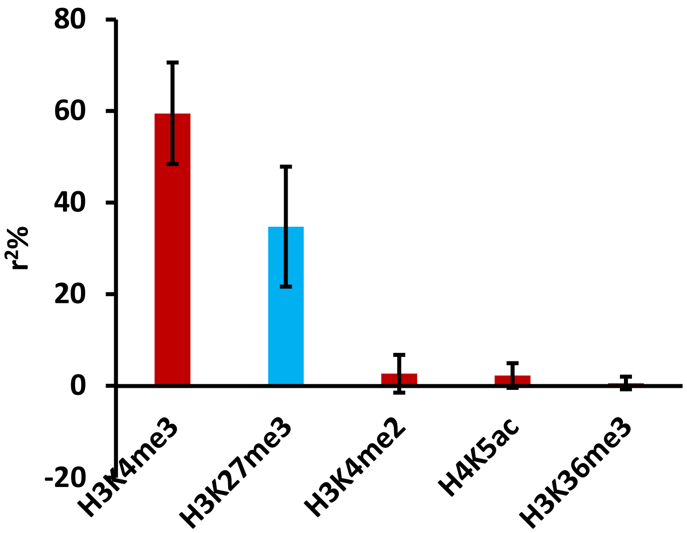


Figure S5. Multiple linear regression and best model selection reveal the predominant epigenetic marks that explain the expression levels of genes associated with energy metabolism.
A. The performance of models generated by different combinations of epigenetic marks. B. Bayesian Information Criterion (BIC) reveals the relationship between model complexity and prediction accuracy. The red dot represents the number of variables to achieve the least complex and most accurate model. Based on adjusted $R^2$ and BIC, the model containing 5 variables was selected as the best one to predict the expression levels of genes associated with energy metabolism. C. Bar plot represents the relative contribution of each epigenetic mark in the selected model containing 5 variables. The bars were color coded based on the role of each epigenetic mark on gene expression. Red represents activation marks and blue represents repression marks.

A.

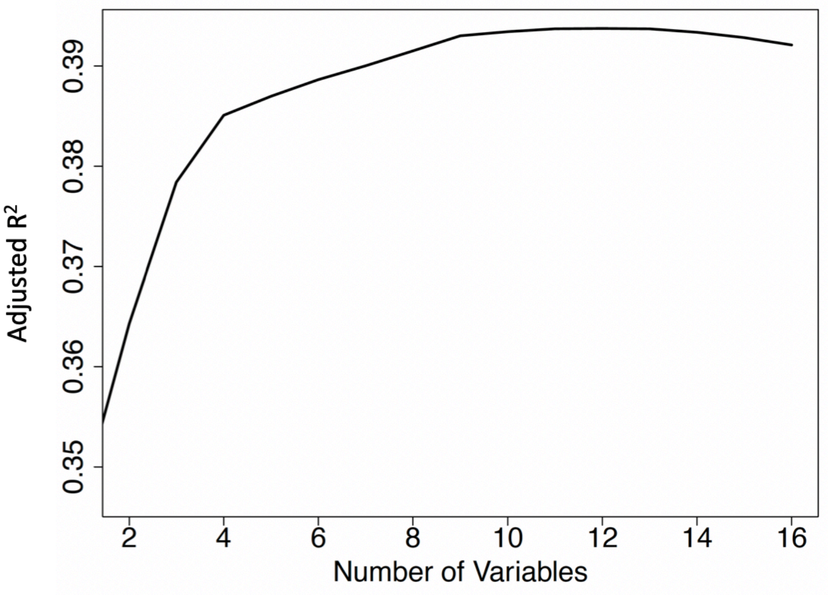


B.

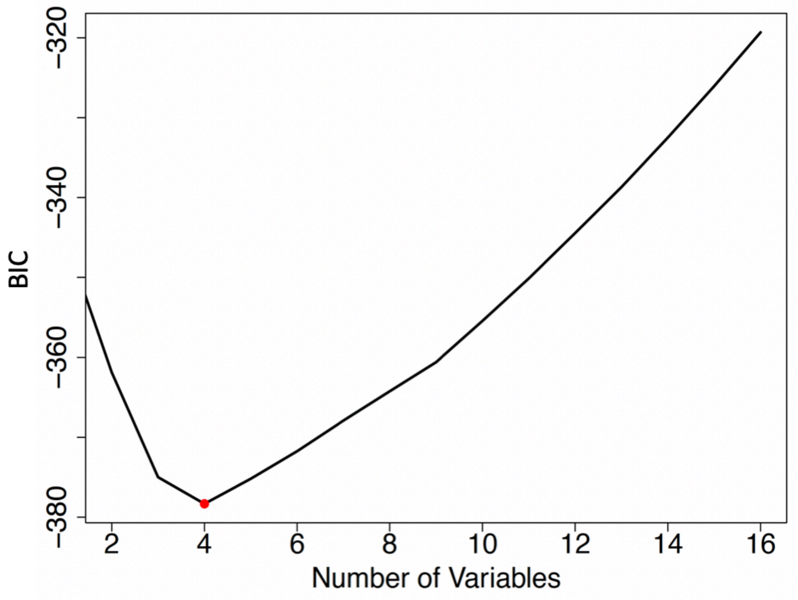


Figure S6. Multiple linear regression and best model selection reveal the predominant epigenetic marks that explain the expression levels of specialized metabolism-related genes. A. The performance of models generated by different combinations of epigenetic marks. B. Bayesian Information Criterion (BIC) reveals the relationship between model complexity and prediction accuracy. The red dot represents the number of variables to achieve the least complex and most accurate model. Based on adjusted $R^2$ and BIC, the model containing 4 variables was selected as the best one to predict the expression levels of genes associated with specialized metabolism.

A.

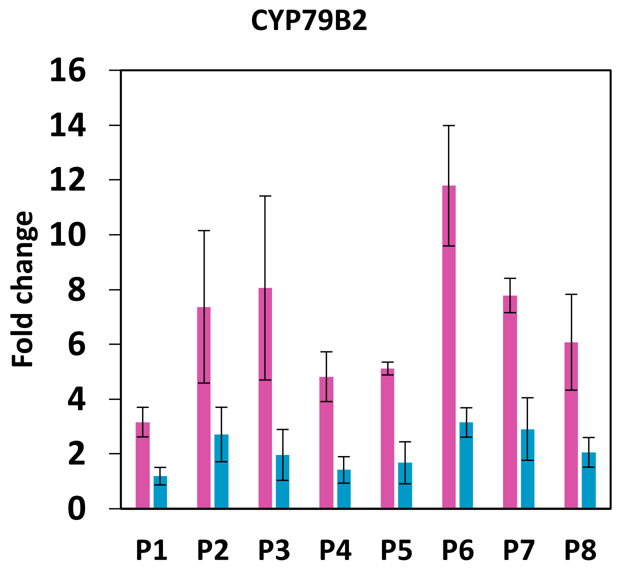
CYP79B2
16
14
12
10
8
6
4
2
0
Fold change
P1
P2
P3
P4
P5
P6
P7
P8

B.

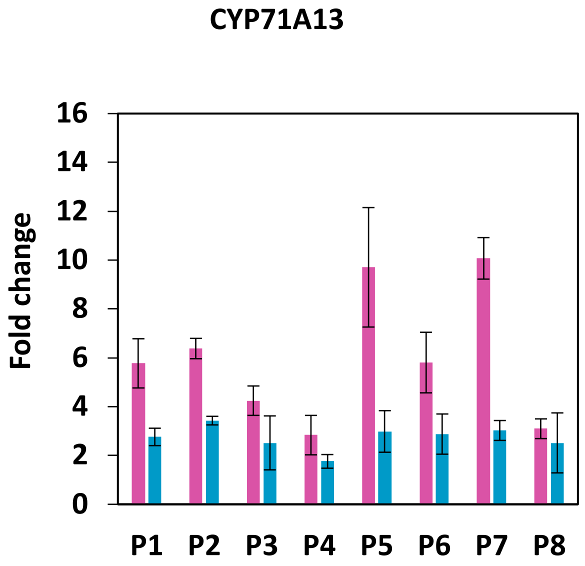
CYP71A13
16
14
12
10
8
6
4
2
0
Fold change
P1
P2
P3
P4
P5
P6
P7
P8

C.

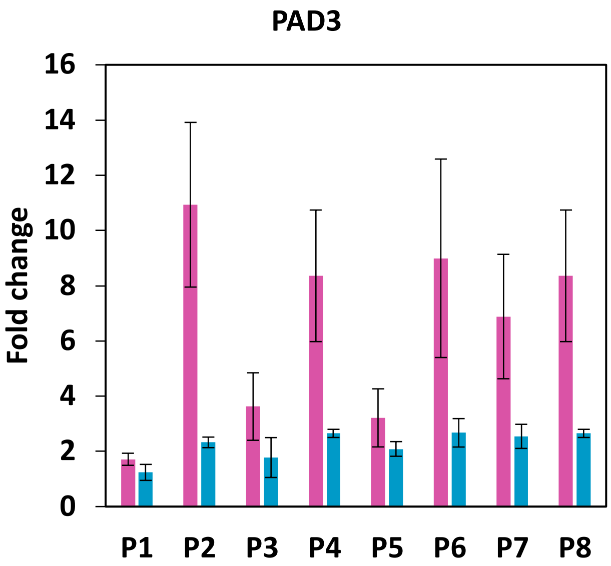


Figure S7. ChIP-qPCR shows that the abundance of H3K27me3 is significantly reduced in the regions of camalexin biosynthesis genes in *pkl-1* mutant than that in wild type plants. P1 to P8 represent the genomic regions that were examined using ChIP-qPCR, which covered 1kb upstream and the transcript region of each gene A. B. and C. represent the ChIP-qPCR results for the three genes in camalexin biosynthesis pathway. Fold change was calculated by normalizing the ChIP signal to that in the negative control assays without H3K27me3 antibody. Error bars represent standard deviations of three biological replicates. Purple bars represent Col-0 and blue bars represent *pkl-1*.

A.

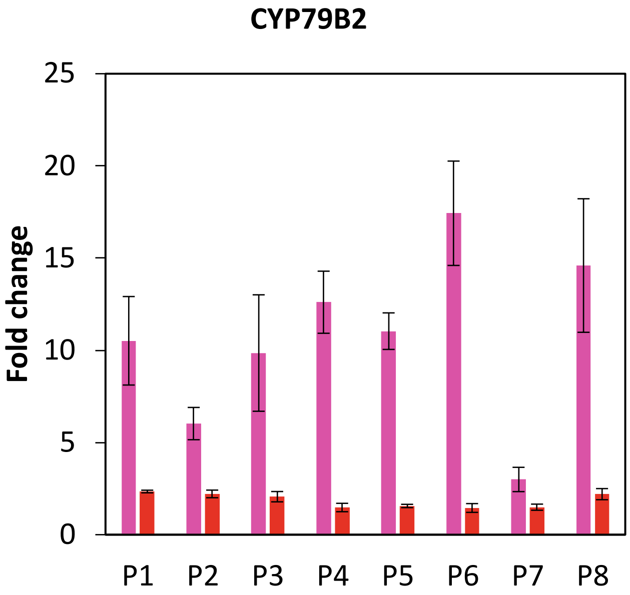
CYP79B2
Fold change
0
5
10
15
20
25
P1
P2
P3
P4
P5
P6
P7
P8

B.

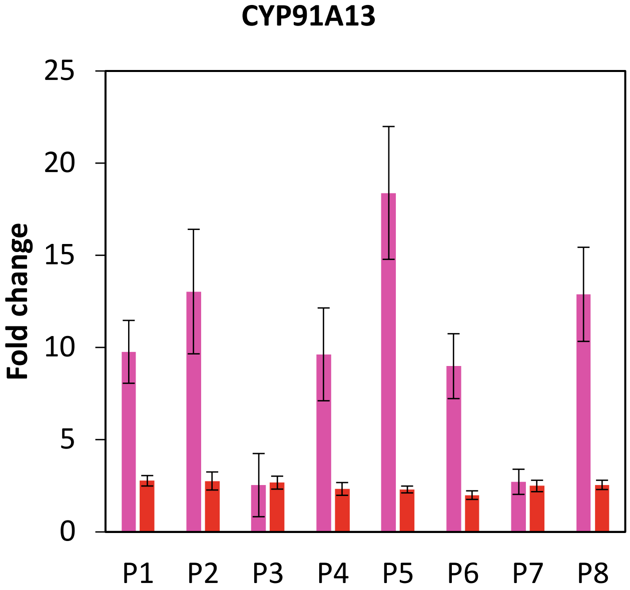
CYP91A13
Fold change
0
5
10
15
20
25
P1
P2
P3
P4
P5
P6
P7
P8

C.

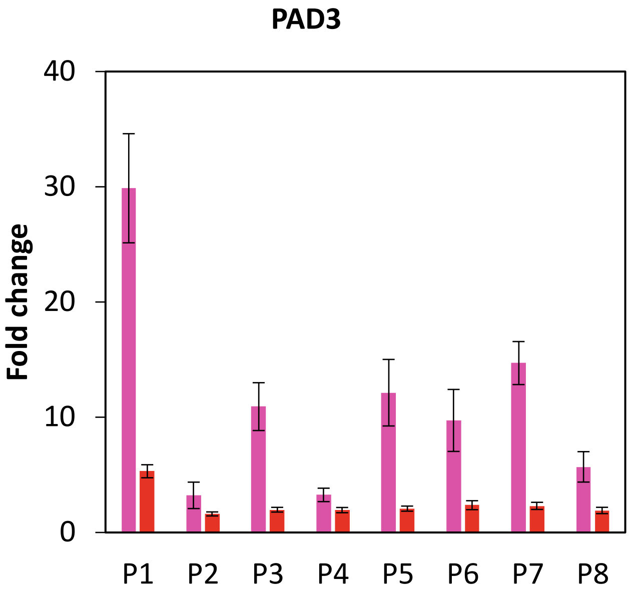


Figure S8. ChIP-qPCR shows that the abundance of H3K18ac is significantly reduced the regions of camalexin biosynthesis genes in *had18-1* mutant than that in wild type plants. P1 to P8 represent the genomic regions that were examined using ChIP-qPCR, which covered 1kb upstream and the transcript region of each gene. A. B. and C. represent the ChIP-qPCR results for the three genes in camalexin biosynthesis pathway. Fold change was calculated by normalizing the ChIP signal to that in the negative control assays without H3K18ac antibody. Error bars represent standard deviations of three biological replicates. Purple bars represent Col-0 and red bars represent *hda18-1*.

**Table S1**. Primers used to quantify the abundance of H3K27me3 and H3K18ac in the genomic regions of camalexin biosynthesis genes using ChIP-qPCR in wild type Col-0, *pkl-1*, and *hda18-1* plants.

| Gene | Primer | Sequence |
|---|---|---|
| CYP79B2 | P1-F | GACTTAAGACTTAGTCTCGGT |
| CYP79B2 | P1-R | TAGCCATTATGTTTTGGGTACG |
| CYP79B2 | P2-F | CAATGTATACGTTACAGAT |
| CYP79B2 | P2-R | TTATGTCTAGCTAGTATTG |
| CYP79B2 | P3-F | CGAAGTATTAGATCAATGT |
| CYP79B2 | P3-R | TCCTCGTGTTATATATGCAC |
| CYP79B2 | P4-F | GGCAGGTCACCAACAAAAC |
| CYP79B2 | P4-R | GTAGCATCACTAAGGTTATAG |
| CYP79B2 | P5-F | CAAGAAATTGATGACGGATC |
| CYP79B2 | P5-R | AGAGAGGATCTTCTGAGCG |
| CYP79B2 | P6-F | GCGTGATCACTCCCTTTGGT |
| CYP79B2 | P6-R | TCCACCGTCAGGTGCAGTG |
| CYP79B2 | P7-F | CCCACCGTAGAAGATGTAGA |
| CYP79B2 | P7-R | TGCCTTGTTCGTCTTTGAT |
| CYP79B2 | P8-F | GCTTACCGCCGATGAAATCA |
| CYP79B2 | P8-R | TGACTTCCTTTAGGGATGTG |
| CYP71A13 | P1-F | GTTCATCATCACTAGTCTTAC |
| CYP71A13 | P1-R | CATAAGTCTTAAGATCGACG |
| CYP71A13 | P2-F | GCAGGATTTACTGAGTTAAAG |
| CYP71A13 | P2-R | GGAATTAAACGTAATCTTTC |
| CYP71A13 | P3-F | CTACTATCATAGGTCGGCT |
| CYP71A13 | P3-R | GGAAGAGTTTATTAGCGACA |
| CYP71A13 | P4-F | TAGCATGCAGAATATGAGT |
| CYP71A13 | P4-R | GAACGGTGAGGATGGAGGC |
| CYP71A13 | P5-F | CTCAGTCTCAGGTACGGAC |
| CYP71A13 | P5-R | CACTCTTCATCTGTCTCCAG |
| CYP71A13 | P6-F | TGGTTGAATCCTTTGAGAAG |
| CYP71A13 | P6-R | GCCTCACTCGCTTCTTGAG |
| CYP71A13 | P7-F | CATATTGGCATGGATAGATG |
| CYP71A13 | P7-R | GTAGAGTCGAAGTTGTTGACG |
| CYP71A13 | P8-F | GGACGATGACGGAACTGATC |
| CYP71A13 | P8-R | GCAGTGTCTCGTTGGATCG |
| PAD3 | P1-F | CACCGCTAAAATTGTTGAC |
| PAD3 | P1-R | CATGTTTATCCTTAATGAGC |
| PAD3 | P2-F | GGATTTGTAATCTACTTTC |

| | | |
|---|---|---|
| PAD3 | P2-R | CAGTATTGGGTCGTCTCAAGC |
| PAD3 | P3-F | CGGTCAGTGAAGTCTACAT |
| PAD3 | P3-R | GAATAATAATCGTAAGTGGAC |
| PAD3 | P4-F | CAAGCTACAGCGGATAGTAG |
| PAD3 | P4-R | TTGGACCCGGAGGAAGCTTA |
| PAD3 | P5-F | AGAAGCTTCCCATCATCGG |
| PAD3 | P5-R | CATCCCGATGTCTTTGAAG |
| PAD3 | P6-F | CGGTGACGAGTGGAGTCTGA |
| PAD3 | P6-R | TCCACGACTCTATCAGCTTC |
| PAD3 | P7-F | GTGCTAAAGGCTGAAGCGGT |
| PAD3 | P7-R | GTGGTGAACTTGAGAGCATC |
| PAD3 | P8-F | CGGACATATTTGTAGCAGG |
| PAD3 | P8-R | AAGAGTGGAGTTGTTGGATG |

**Table S2**. Primers used in the sequential ChIP-qPCR experiment to examine the co-localization of H3K27me3 and H3K18ac within camalexin biosynthesis genes.

| Name | Sequence |
|---|---|
| CYP79B2_qF | GCAATGGAAGAGATCGACAGAG |
| CYP79B2_qR | GGAGGATAGCTTTGACGTAGTTTAG |
| CYP71A13_qF | CCAACGAGACACTGCGATATG |
| CYP71A13_qR | GATCCGAATGGGATGTAGTTCAG |
| PAD3_qF | GCAGCAGAGGAAGTGCTAAA |
| PAD3_qR | ATCCCGATGTCTTTGAAGTTGT |
| GLK1_qF | GATTTAGAGCACCGCCAGTT |
| GLK1_qR | GAGCACCACCAAATCCAAGA |
| H3K27me3_only_qF | CAACGGTTCTTCATCCGATT |
| H3K27me3_only_qR | CTGCTCGAAATGGCTCTACC |
| H3K18ac_only_qF | GGTAACCGATGTGGGACATTT |
| H3K18ac_only_qR | CCAACAAGACTGGTCCAAAGA |
| Negative_con_qF | TGCTCGTCCCATTTCCTATC |
| Negative_con_qR | GGCATAGTGATTTTGCCACA |

**Table S3**. Primers used to examine the expression of camalexin biosynthesis genes under FLG22 induction in wild type Col-0, *pkl-1*, and *hda18-1* plants.

| Name | Sequence |
|---|---|
| CYP79B2_qF | ATCTCCTCTCAACACTTCAAGC |
| CYP79B2_qR | GGTGGCAGATACGGTTTCTTT |
| CYP71A13_qF | CGATTTGACTGGAGGGTAGAG |
| CYP71A13_qR | CCGAAGATGGAAATGCAATGAG |
| PAD3_qF | GCAGCAGAGGAAGTGCTAAA |
| PAD3_qR | ATCCCGATGTCTTTGAAGTTGT |

**Data S1**. $Log_2$(fold change) in the enrichment analysis to identify predominant epigenetic modifications in energy metabolism-related pathways. Only the pathways that contain at least ten genes were included in this analysis.

**Data S2**. $Log_2$(fold change) in the enrichment analysis to identify predominant epigenetic modifications in specialized metabolism-related pathways. Only the pathways that contain at least ten genes were included in this analysis.
